# Making Aptamers More Antibody-like: Targeting AXL *in Vivo* Using a Bottlebrush Polymer-Conjugated Aptamer

**DOI:** 10.1101/2024.10.02.616316

**Authors:** Tingyu Sun, Jiachen Lin, Chenyang Xue, Yuyan Wang, Peiru Chen, Yun Wei, Guobin Xu, Anais Sidonia, Chris Nenopoulos, Hossam Tashkandi, Ke Zhang

**Affiliations:** Department of Chemistry and Chemical Biology, Northeastern University, Boston Massachusetts, 02115, USA; Department of Bioengineering, Northeastern University, Boston, Massachusetts, 02115, USA; Department of Chemical Engineering, Northeastern University, Boston, Massachusetts, 02115, USA; Department of Biochemistry, Northeastern University, Boston, Massachusetts, 02115, USA

**Keywords:** Oligonucleotide, aptamer, AXL, drug resistance, cancer

## Abstract

The overexpression of receptor tyrosine kinase AXL is linked to acquired drug resistance in cancer treatments. Aptamers, acting as antibody surrogates, have been envisioned as potential inhibitors for AXL. However, aptamers face difficult pharmacological challenges including rapid degradation and clearance. Herein, we report a phosphodiester-backboned bottlebrush polymer as a carrier for conjugated aptamers. Termed pacDNA, the conjugate improves aptamer specificity *in vivo*, prolongs blood retention, and enhances overall aptamer bioactivity. Treatment with pacDNA in AXL-overexpressing cell lines significantly inhibits AXL phosphorylation, resulting in reduced cancer cell migration and invasion. In a non-small cell lung cancer xenograft model (NCI-H1299), pacDNA treatment leads to single-agent reduction in tumor growth. These results highlight the potential of bottlebrush polymers in the field of aptamer therapeutics.

## Introduction

AXL is a member of the TAM (TYRO3-AXL-MER) receptor tyrosine kinase subfamily.^1^ Upon binding with its primary ligand, growth arrest-specific 6 (GAS6) protein, AXL undergoes dimerization and autophosphorylation of tyrosine residues, initiating downstream signaling cascades involved in various cellular processes such as survival, growth, differentiation, adhesion, proliferation, and invasion.^2–6^ In healthy adults, AXL expression is typically low; however, aberrant upregulation of GAS6/AXL has been observed in many human malignancies, showing correlation with tumor resistance to chemotherapy, programmed death-1 (PD-1) inhibitors, targeted therapies, and radiation therapy.^7–10^ AXL-targeted approaches have shown promise in overcoming drug resistance, with several therapies currently in the clinic, including small molecule selective inhibitors, multitargeted inhibitors, antibody-drug conjugates, and anti-AXL Fc fusion proteins.^11–13^ However, there are currently no FDA-approved drugs targeting AXL.^14^

Aptamers, synthetic single-stranded oligonucleotides with specific tertiary structural interactions, have emerged as a promising modality to target AXL.^15^ Aptamers are free from potential toxicities arising from unintended kinase inhibition by small molecule inhibitors or immunogenic side effects of biologics, and are versatile with regard to conjugation approaches.^16,17^ A modified DNA aptamer targeting AXL has been reported, which showed potent antitumor effects in *in vitro* and *in vivo*.^18^ Nonetheless, aptamers as a modality still face decades-long challenges of rapid glomerular clearance and potential cross-reactivity outside of the ideal environment in which the aptamer was selected.^19,20^

Here, we demonstrate that a bottlebrush polymer can be used to enhance the bioactivity of an AXL-targeted aptamer *in vivo*. The bottlebrush polymer consists of a poly(serinol phosphodiester) (PSP) backbone and 30 poly(ethylene glycol) (PEG) side chains covalently conjugated to the backbone.^21^ A single aptamer is attached to the terminus of the backbone. Termed pacDNA (polymer-augmented conjugates of DNA), the conjugate creates a spatially congested PEG environment, which mitigates aptamer nonspecific binding, while the large size (∼320 kDa) of the conjugate limits renal clearance. Together, these properties result in prolonged blood retention times, enhanced specific binding *in vivo*, and potent single-agent anti-tumor activity in an NCI-H1299 xenograft mouse model. Overall, the pacDNA system we established here offers a promising approach as a universal platform to impart aptamers with antibody-like *in vivo* properties.

**Scheme 1.**
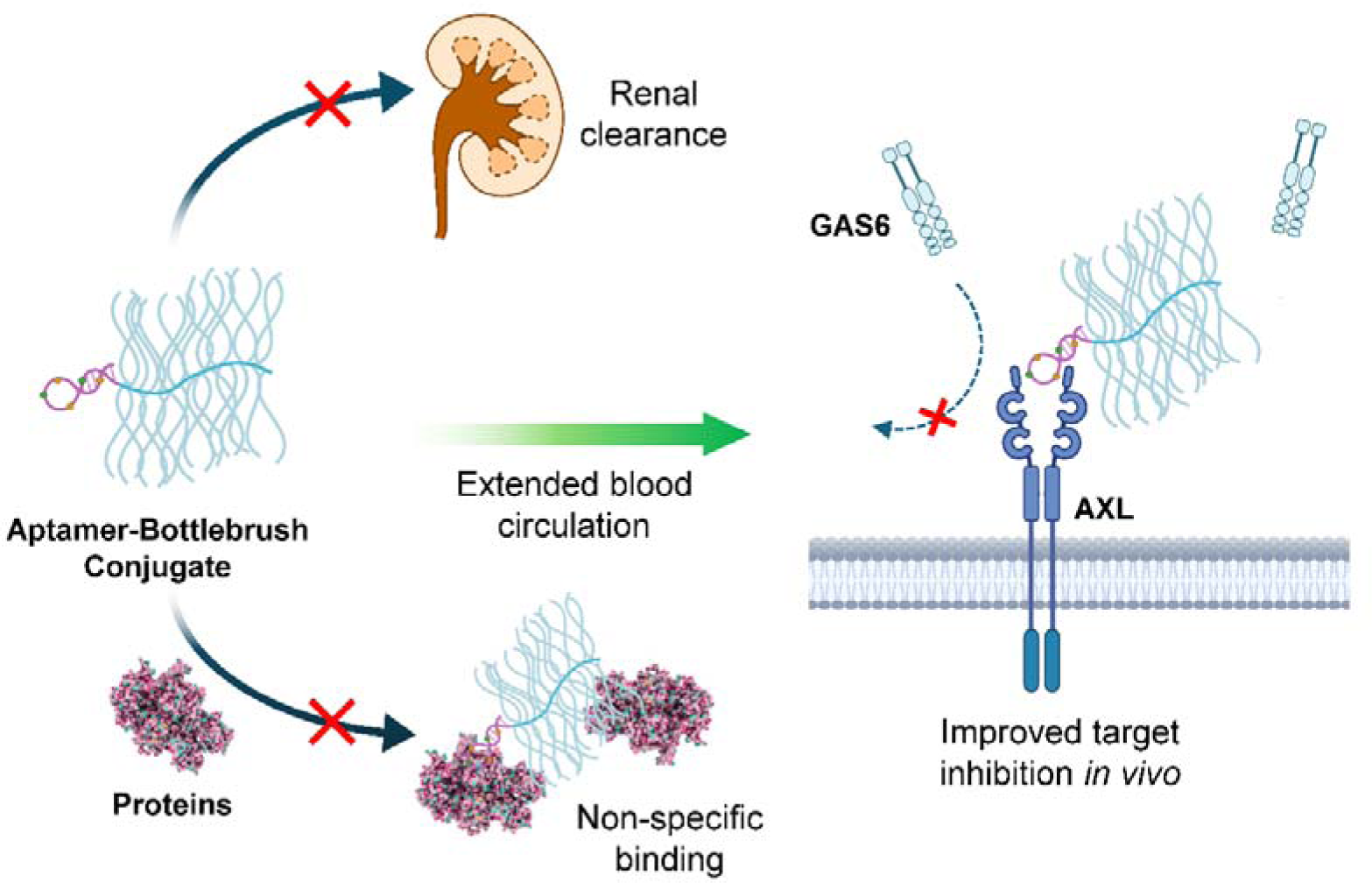
Schematic illustration showing pacDNA’s improved *in vivo* characteristics and selective inhibition of AXL signaling.

## Result and Discussion

### Preparation and Characterization of PSP pacDNA

The construction of the bottlebrush polymer PSP backbone involves a sequential condensation process using a bespoke Fmoc-protected phosphoramidite (Figure 1A). Because this synthesis shares the same phosphoramidite chemistry as solid-phase oligonucleotide synthesis, the aptamer component can be integrated into the polymer backbone in one synthesis, eliminating the necessity for a post-conjugation step.^21^ A PSP backbone comprising 30 repeating units and one AXL-binding aptamer positioned at its 3’ end was synthesized. The aptamer sequence (Table S1) was modified from GLB-A04 (replacement of phosporodithoate [PS2] with phosphorothioate [PS]), which was reported by Lopez-Berestein.^18^ Following the synthesis, the Fmoc protective groups on the serinol amines were removed, and purification was performed using reversed-phase high-performance liquid chromatography (RP-HPLC, Figure S1). Subsequently, the backbone strands underwent two-stage PEGylation (first in aqueous buffer then in dimethyl formamide) and removal of excess PEG by aqueous gel permeation chromatography (GPC), resulting in high molecular weight (M_n_: 301 kDa) and low polydispersity (PDI: 1.2, Figure S3) bottlebrush conjugates (pacDNA, Figure 1c) with high batch-to-batch consistency. Transmission electron microscopy (TEM, Figure 1B and Figure S2) confirmed that these molecular brushes exhibit a spherical morphology, non-aggregation, and size uniformity when dispersed in water, which is corroborated by dynamic light scattering (DLS, Figure S4), showing monomodal distribution and a hydrodynamic diameter of 27.7 ± 8.6 nm (intensity average).

**Figure 1.**
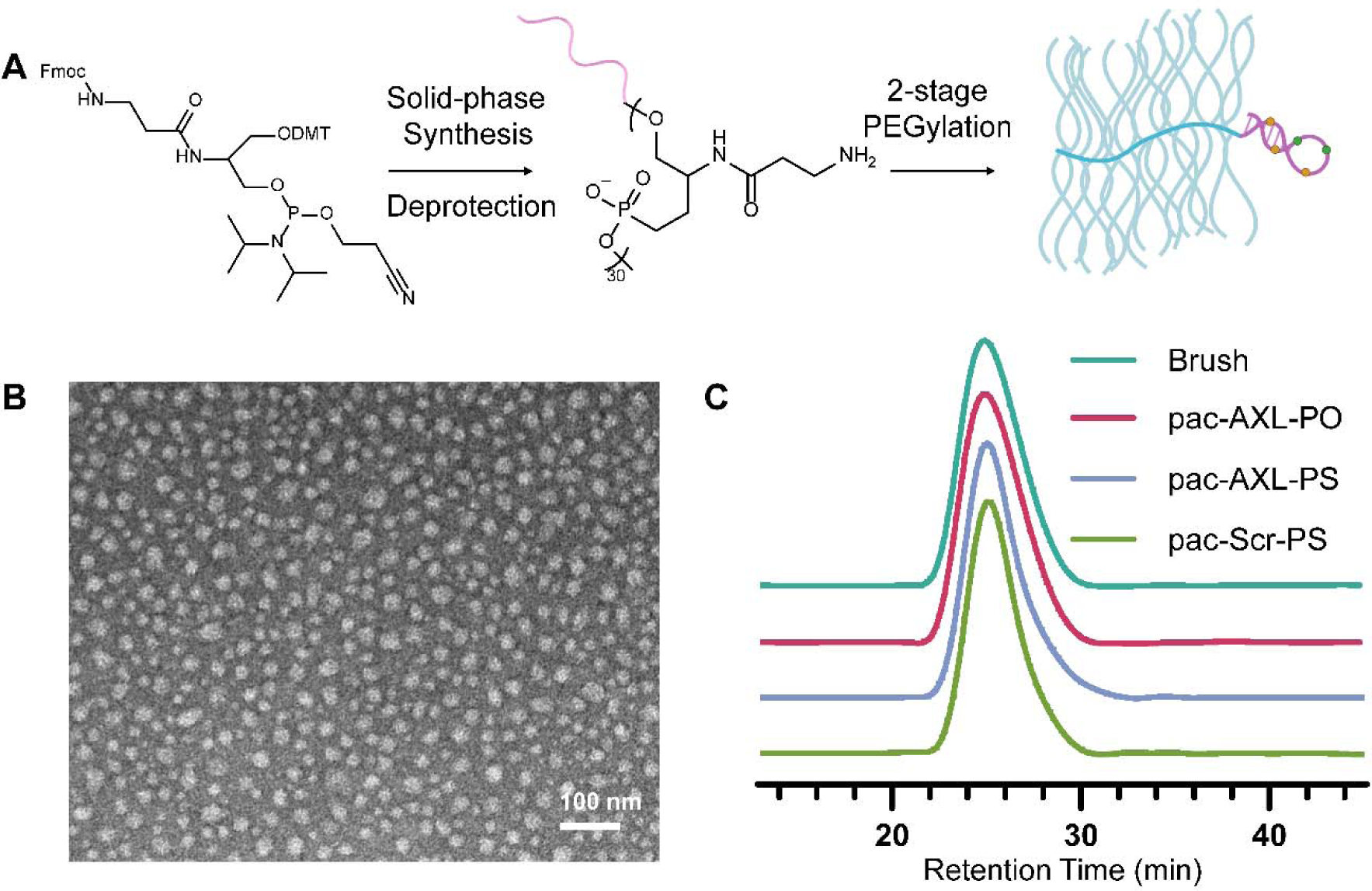
Characterization of PSP pacDNA. (A) Schematics of the synthesis of PSP bottlebrush polymer-aptamer conjugates. (B) Representative TEM image of pac-AXL-PS. (C) Aqueous GPC chromatograms of the PSP bottlebrush polymer and aptamer/control conjugates.

### Binding affinity analysis

Previous docking studies using Autodock Vina demonstrated that GLB-A04 has affinity for the extracellular domain (ECD) of the AXL receptor, which consists of two immunoglobulin-like (Ig1 and Ig2) domains and two fibronectin type 3-like domains.^18^ The stem region of GLB-A04 interacts with the AB loops of the Ig1 domains, with a binding affinity of 592 ± 92 nM. We modified the GLB-A04 aptamer by replacing the PS2 linkage between the first two nucleotides at the 5’ with a PS due to the poor yield of the desulfurization. Interestingly, when the binding affinity of the modified PS aptamer (AXL-PS) and its corresponding pacDNA (pac-AXL-PS) was evaluated by microscale thermophoresis (MST) utilizing Cy5-labeled aptamers as the detection agent, it was discovered that AXL-PS exhibits improved binding (84 ± 12 nM) to AXL (Figure 2C). A modest decrease in affinity was observed upon integration into the bottlebrush structure, with the dissociation constants for pac-AXL-PS determined to be 163 ± 26 nM. Importantly, two control pacDNAs, one with PS chemistry but scrambled nucleotides (pac-Scr-PS) and the other with the correct nucleotide sequence but no chemical modifications (full phosphodiester [PO], pac-AXL-PO), demonstrated minimal interactions with AXL, confirming specific binding rather than non-specific interactions with the bottlebrush component. Further investigations into the specificity of aptamer interactions *in vitro* were conducted by incubating the pacDNAs and controls with four distinct cancer cell lines with varying endogenous levels of AXL expression (Figure S5) and subsequent analyses of cell-associated fluorescence using flow cytometry (Figure 2A). Cells were pre-treated with 0.1% sodium azide solution for 1 h to deplete adenosine triphosphate (ATP) and suppress cell endocytosis, limiting signals to cell surface-bound materials. It was revealed that cell-associated signals increased with AXL expression levels with significant linearity (R^2^>0.87) for AXL-PS and pac-AXL-PS, while negative controls including Scr-PS, pac-Scr-PS, and pac-AXL-PO exhibited little to no AXL-dependent cell binding (Figure 2B). Further, when extrapolating the linear fit of cell-bound signals to zero on the x-axis (relative AXL expression levels), there is a significant intercept with the y-axis for AXL-PS, suggesting that these PS oligonucleotides non-specifically bind to cell surface proteins in the absence of AXL. The non-specifically bound materials can contribute to a significant portion of the total cell-associated materials. For example, for SKOV3 cells which express a low level of AXL, 79% of cell-bound materials is estimated to be non-specific to AXL. Such unwanted interactions may be particularly detrimental for *in vivo* applications of PS aptamers, where the aptamer must circulate for a sufficiently long time to reach its intended receptors. A background level of non-specific binding of the PS oligonucleotide with non-targeted cells and tissues may rapidly deplete them from blood circulation, resulting in suboptimal targeting. On the other hand, the linear fit of pac-AXL-PS shows minimal y-intercept, suggesting that the bottlebrush polymer reduces non-specific binding. Indeed, previous studies have shown that a pacDNA containing double-stranded DNA is more resistant to nuclease degradation than free DNA, which is attributed to an entropic effect created by the dense arrangement of the side chains.^22^ This effect broadly reduces oligonucleotide-protein interactions, leading to a corresponding reduction in side effects that derive from these interactions, such as coagulopathy and unwanted activation of the immune system.^23–26^

**Figure 2.**
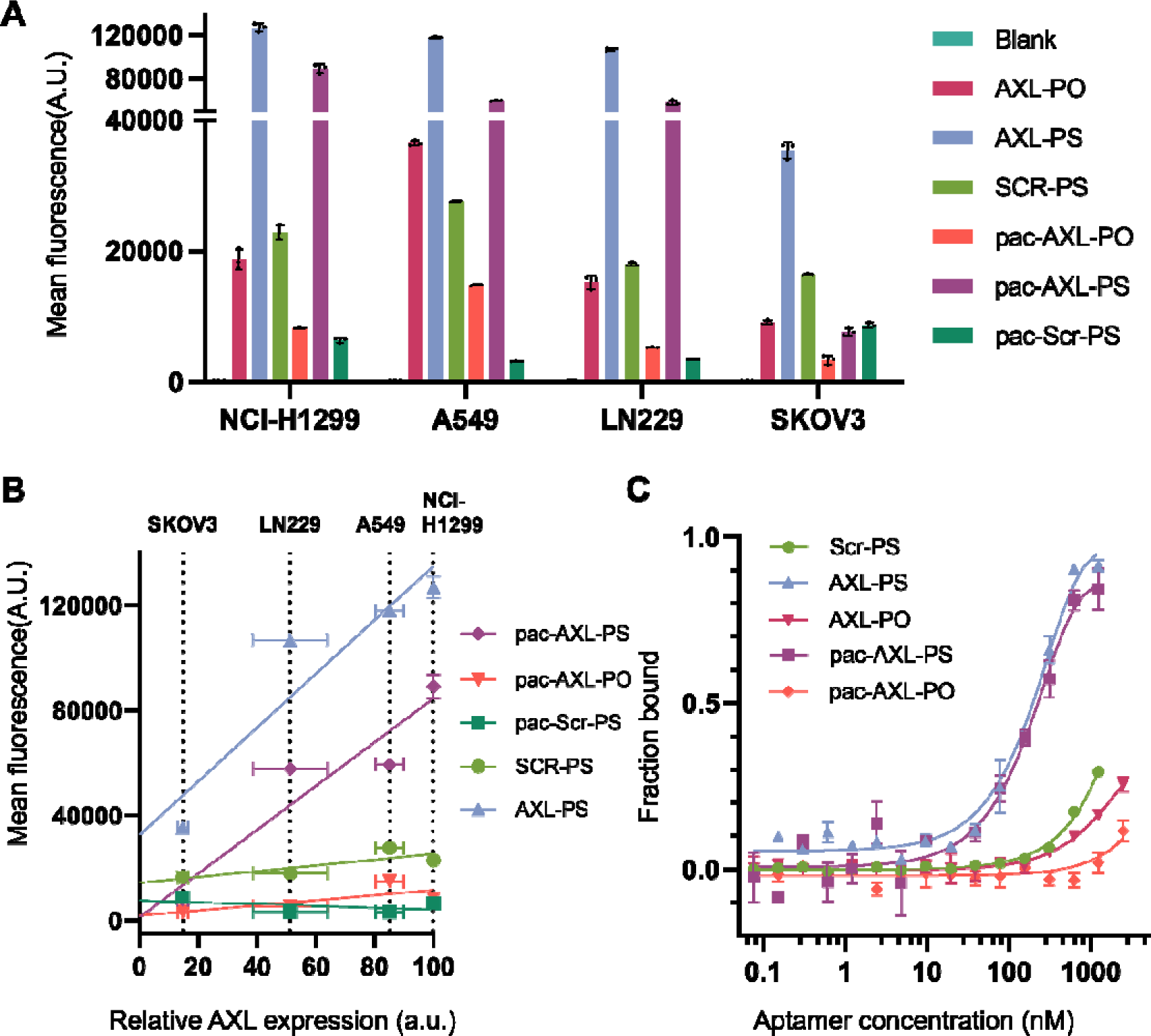
Binding affinity analysis. (A) Cell-associated materials (NCI-H1299, A549, LN229, and SKOV3 cells) after incubation with samples and controls as determined by flow cytometry. (B) Cell-associated materials as a function of endogenous AXL expression levels. (C) Binding of samples and controls with isolated human AXL protein as measured by microscale thermophoresis.

### *In vitro* functional study

To investigate the functional activity of pac-AXL-PS on AXL phosphorylation, three AXL-overexpressing cell lines (NCI-H1299, LN229, and A549) were treated with samples and controls at concentrations ranging from 0.1 to 5 μ (aptamer basis) in the presence of GAS6, which stimulates AXL phosphorylation. Immunoblotting analysis of cell lysates indicated concentration-dependent reduction of AXL phosphorylation for cells treated with AXL-PS and pac-AXL-PS. In contrast, Scr-PS and pac-Scr-PS showed minimal effect on AXL phosphorylation (Figure 3A). Interestingly, pac-AXL-PS exhibited more potent reduction of AXL phosphorylation than AXL-PS. One interpretation is that pac-AXL-PS can sterically enhance the antagonistic effect of the aptamer against competitive ligands (*e.g.* GAS6) due to the large size of the bottlebrush polymer.

**Figure 3.**
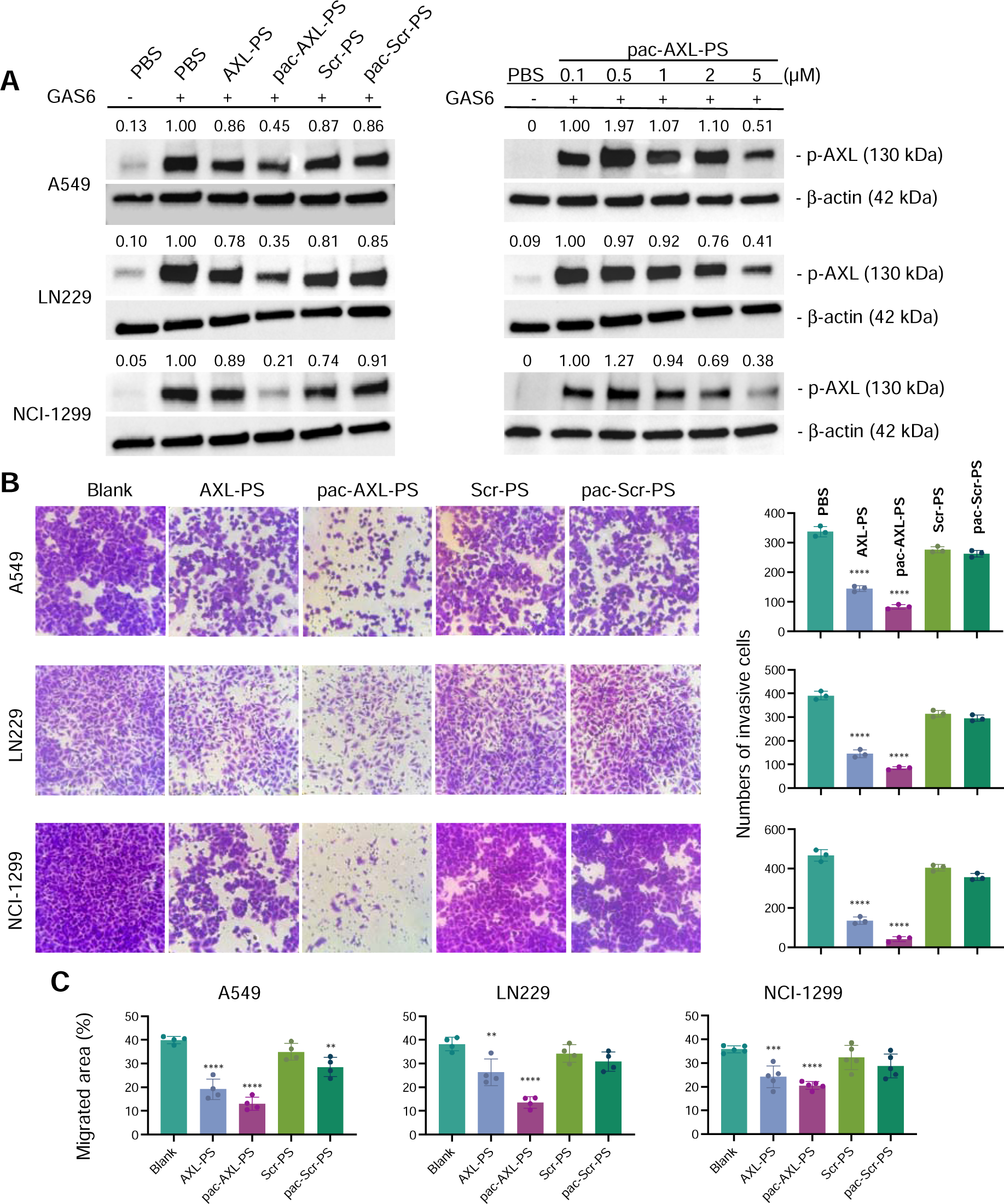
*In vitro* functional studies of aptamers and pacDNA. (A) Inhibition of AXL phosphorylation by aptamers and pacDNA in A549, LN229, NCI-H1299 cells (left panel) and concentration dependence for pac-AXL-PS (right panel). (B) Invasion inhibition in A549, LN229, and NCI-H1299 cells (left panel) and quantification of invading cells by cell counting (right panel). (C) Inhibition of cell migration as determined by the cell migration scratch assay. Images used for the analysis can be found in Figure S6. Statistical significance was calculated using Student’s two-tailed t-test. ***P<0.001, ****P<0.0001

We next probed the aptamer’s downstream impact on metastasis, specifically cellular invasion and migration. Cell invasiveness (Figure 3B) was assayed using a transwell chamber coated with Matrigel. NCI-H1299, LN229, and A549 cells, suspended in serum-free medium containing samples or controls, were added to the upper chambers, while medium with 10% fetal bovine serum was loaded in the lower chamber serving as a chemo-attractant. After the incubation, cells that migrate through the Matrigel are considered invading cells, which are stained with 0.1% crystal violet for counting. Treatment with pac-AXL-PS resulted in a marked decrease of invading cells. The fraction of invading cells from the corresponding non-treated group for A549, LN229, and NCI-H1299 cells are 24%, 21%, and 9%, respectively. Again, pac-AXL-PS appears to be superior to AXL-PS alone, consistent with *in vitro* AXL phosphorylation results (*vide supra*). The anti-invasion effect appears to be specific to AXL binding, as both Scr-PS and pac-Scr-PS exhibited no significant inhibition. To measure cell migration (Figure 3C and supplementary Figure S6), cells were cultured until reaching 100% confluence to form monolayers. Each monolayer was scraped with a pipette tip and subsequently treated with samples (2 µM equiv. of aptamer) or controls. Cell migration was quantified by measuring the area not covered by cells from the scrape edge after 24 hours. Treatment with pac-AXL-PS showed the most significant inhibition of cell migration following the 24-hour period across all examined cell lines, whereas Scr-PS and pac-Scr-PS did not exhibit notable inhibitory effect.

### Plasma pharmacokinetics and antitumor efficacy *in vivo*

To evaluate the plasma pharmacokinetics (PK) of the aptamer, blood samples were collected from C57BL/6 mice following intravenous (i.v.) administration of Cy5-labeled samples and controls (Figure 4B). Free aptamers were swiftly eliminated from circulation, likely via renal glomerular filtration, resulting in very short half-lives (0.48 h for AXL-PS and 0.14 h for AXL-PO, t_1/2α_, two-component model). In contrast, both pac-AXL-PO and pac-AXL-PS exhibited α significantly prolonged circulation times, showing two orders of magnitude greater area under the curve (AUC) compared to AXL-PO (Table S2). The unconjugated polymer, being a stealth platform, displays the highest level of blood retention with approximately 20% of the injected dose still in circulation 72 h post injection. These findings indicate that enzymatic and chemical stability of the aptamer are secondary contributing factors to PK. The bottlebrush polymer plays a dominating role in blood concentration and bioavailability.

**Figure 4.**
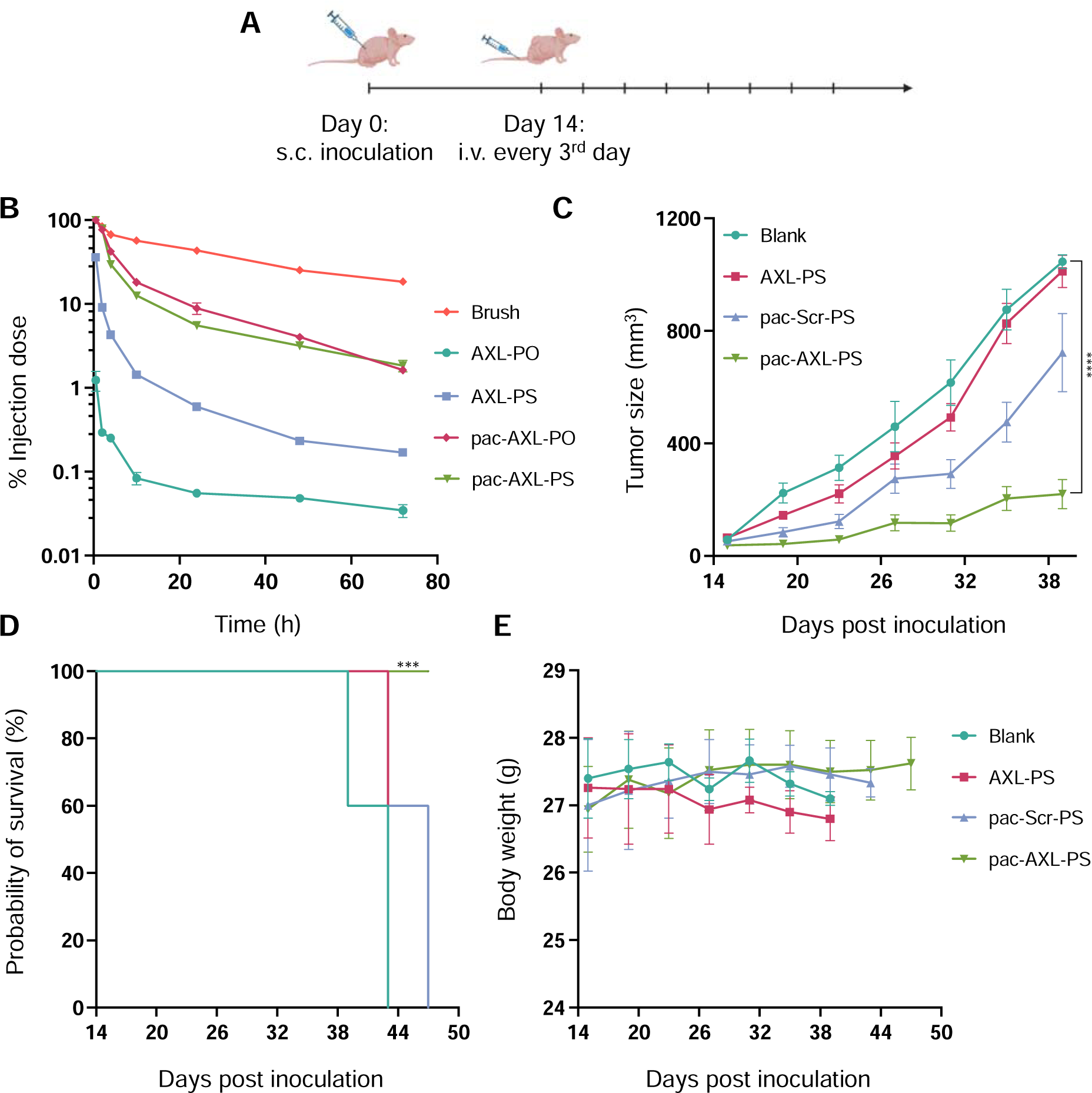
*In vitro* efficacy studies of aptamers and pacDNA. (A) Tumor inoculation and treatment schedule. (B) Plasma pharmacokinetics of samples and controls. (C) Average tumor growth curve after mice were inoculated with NCI-H1299 cells. (D) Kaplan-Meier animal survival analysis. (E) Animal body weight changes during the treatment period. Statistical significance was calculated using two-way ANOVA. ****P<0.0001.

The efficacy of pac-AXL-PS in inhibiting tumor growth was evaluated in female BALB/c nu/nu mice with subcutaneous NCI-H1299 xenografts. When the xenografts reached approximately 80 mm^3^ in volume, pac-AXL-PS, free AXL-PS, or controls were administered i.v. (0.5 μmol/kg) once every third day for a total of 8 doses (Figure 4A). By day 38, the average tumor volume in the saline-treated groups had progressed to around 900 mm^3^. Notably, pac-AXL-PS displayed potent single-agent tumor growth inhibition (102-336 mm^3^; Figure 4C). In contrast, while AXL-PS displayed a measurable level of bioactivity *in vitro*, it exhibited negligible antitumor effect *in vivo*. This discrepancy can be attributed to the pharmacokinetic challenges associated with unformulated aptamers, which must overcome strong competing factors including renal clearance and adsorption by non-tumor tissues, resulting in much subsided responses *in vivo*. Studies carried out under ideal *in vitro* conditions do not replicate these pharmacokinetic factors. To exclude nonspecific antitumor activity stemming from the PS modification or the polymer component, pac-Scr-PS was employed as a control, which yielded insignificant antitumor responses (Figure 4C and S7). Kaplan–Meier survival analysis (utilizing a fourfold increase in tumor size as a surrogate for survival endpoint; Figure 4D) indicates that treatment with pac-AXL-PS delays the time to reach the surrogate endpoint compared to the control groups. Treatment with pac-AXLs is well-tolerated in mice, as evidenced by the absence of significant body weight loss (Figure 4E) or noticeable changes in behavior (*e.g.*, refusal to eat, startle response).

### Summary

We have demonstrated that conjugation of aptamers with a bottlebrush polymer can effectively overcome the pharmacokinetic difficulties associated with the aptamer and rescue its bioactivity *in vivo*. Using an AXL-targeting aptamer as an example, we show that conjugation with the bottlebrush polymer not only do not interfere with the target binding ability of the aptamer, but also enhances binding specificity and elevates the aptamer’s antagonistic effect *in vitro*, leading to more potent AXL phosphorylation inhibition and cell migration inhibition than the free aptamer. Importantly, the bottlebrush polymer massively improves the blood availability of the aptamer, rendering the aptamer a potent agent for tumor growth inhibition *in vivo*, where the free aptamer failed. These findings underscore the potential of the pac-AXL as a promising agent for addressing the unmet clinical need posed by AXL-driven human cancers. Taken together, the bottlebrush polymer-aptamer conjugate effectively reduces gap between aptamers and antibodies, potentially facilitating aptamer applications in cancer therapeutics and beyond.

## Materials and Methods

### Cell culture

NCI-H1299 cells were cultured in RPMI 1640 media supplemented with 10% fetal bovine serum (FBS) and 1% antibiotics. SKOV3 cells were cultured in DMEM media supplemented with 10% FBS and 1% antibiotics. A549 cells were cultured in F-12K media supplemented with 10% FBS and 1% antibiotics. LN229 cells were cultured in DMEM media supplemented with 5% FBS and 1% antibiotics. All cells were cultured at 37 °C in a humidified atmosphere containing 5% CO_2_.

### Microscale thermophoresis

Microscale thermophoresis (MST) binding measurements were carried out with 10 nM Cy5-labeled samples in binding buffer (1× PBS pH 7.4, 0.1% Triton X 100) with a range of concentration of human AXL protein (154-AL) from 2500 nM to 0.07 nM. The mixtures were transferred to Monolith NT.115 standard capillaries and analyzed on a Monolith NT.115 instrument (NanoTemper Technologies, Munich, Germany) at medium MST power and 20% excitation power. Data was analyzed using the MO. Affinity Analysis software (version 2.3, NanoTemper Technologies) and MST-on time was set at 1.5 s.

### Flow cytometry

Cells were seeded in 24-well plates at a density of 1.0×10^5^ cells per well in 1 mL full growth media and cultured overnight at 37 °C with 5% CO_2_. After incubation, cells were inhibited by 0.1% sodium azide in serum-free culture media for 1 h. After washing by PBS 1×, Cy5-labeled M equiv. of aptamer) dissolved in serum-free culture media (400 µL) were added, and cells were further incubated at 37 °C for 4 h. Next, cells were washed with PBS L per well). Thereafter, 1 mL of PBS was added to each culture μ well to suspend the cells. Cells were then analyzed on a Attune™ NxT flow cytometer (Invitrogen, MA). Data for 1.0×10^4^ gated events were collected.

### Cell migration

5.0 × 10^5^ cells were plated onto six-well plates before treatment with samples (2µM equiv. of aptamer) and controls, and then incubated at 37 °C until 100% confluence is reached. Each cell monolayer was carefully scratched by using a p200 pipet tip, and then cellular debris was rinsed away with 1× PBS buffer. Images (magnification 4×) were captured at 0 and 24 h after scratching using a Nikon eclipse TE2000-U microscope. The percent migration was measured by quantifying the total area that the cells migrated from the scraping edge during 24 h compared to vehicle-treated cells (PBS). All experiments were triplicated.

### Cell invasion

Cell invasision was assessed using a transwell chamber assay. Transwell chambers (Corning, Cellgro) were coated with Matrigel (Corning, Cellgro). NCI-H1299, LN229 and A549 cells suspended in serum-free medium were mixed with samples and controls (2µM equiv. of aptamer) and added to the upper Matrigel-coated chamber (1.0×10^5^ cells/chamber). Cell culture medium with 10% fetal bovine serum was added to the lower chamber as a chemo-attractant. Cells were incubated for 24 h, and the cells in the upper chamber were removed with cotton swabs. The invading cells on the lower surface were fixed with 4% paraformaldehyde and stained using 0.1% crystal violet staining. Cells in three random fields were counted using ImageJ 1.48v software. All experiments were triplicated.

### Immunoblotting

Cells (NCI-H1299, A549, or LN299) were plated at a density of 1.5×10^5^ cells per well in 24-well plates in full medium and cultured overnight at 37 °C with 5% CO_2_. Thereafter, cells were serum-starved for 16 h and then samples and controls (0.1 to 5 μM to the wells and incubated for 0.5 h. After incubation, 400 ng/ml GAS6 (885-GSB) was added to each well for 10 min to stimulate phosphorylation. Then, cells were harvested and lysed in 100 μL of RIPA Cell Lysis Buffer with 1 mM phenylmethanesulfonylfluoride (Cell Signaling Technology, Inc.) following the manufacturer’s protocol. Protein content in the extracts was quantified using a bicinchoninic acid protein assay kit (Thermo Fisher). Equal amounts of proteins (10 μg per lane) were separated on 4 to 20% gradient sodium dodecyl sulfate-polyacrylamide gel electrophoresis and electro-transferred to a nitrocellulose membrane. The membranes were then blocked with 3% bovine serum albumin in Tris-buffered saline with 0.05% Tween-20 and further incubated with appropriate primary antibodies overnight at 4L After washing and incubation with secondary antibodies, proteins were visualized by chemiluminescence using the ECL Western Blotting Substrate (Thermo Scientific). Antibodies used for Western blots were primary anti-p-AXL (1:1000 dilution; D12B2), anti-AXL antibody (1:1000 dilution; AF154), β-actin (1:2000 dilution; AM4302), anti-rabbit IgG, HRP-linked antibody (1:2000 dilution; 7074P2), anti-mouse IgG, HRP-linked antibody (1:5000 dilution; 7076S), and anti-goat IgG, HRP-linked antibody (1:2000 dilution; HAF017). Western blot images were quantified using the Image Lab software by comparing the detected protein band with that of the reference protein.

### Pharmacokinetics

Animal protocols were approved by the Institutional Animal Care and Use Committee of Northeastern University. Animal experiments and operations were conducted following the approved guidelines. 8∼12-week-old female C57BL/6 mice (Charles River, MA, USA) were randomly divided into 5 groups (n=5). Samples and controls were intravenously administered via the tail vein at equal aptamer concentrations (0.5 μmol/kg equiv. of aptamer) were collected from the submandibular vein at varying time points (30 min, 2 h, 4 h, 10 h, 24 h, 48 h, and 72 h) using BD Vacutainer blood collection tubes with sodium heparin. Heparinized plasma samples were obtained by centrifugation at 3000 rpm for 20 min, and then aliquoted into a 96-well plate. The fluorescence intensities were measured on a plate reader. The amounts of agents in the blood samples were estimated using standard curves established for each sample in freshly collected plasma.

### Antitumor efficacy

To establish the NCI-H1299 xenograft tumor model, ∼2×10 cells in 100 μL PBS were implanted subcutaneously on the right flank of 6-week-old BALB/c nude mice. Mice were monitored for tumor growth every two days. When the xenograft reached a volume of 80 mm3, mice were randomly divided into 4 groups (n = 5) to receive the following treatments by intravenous administration: 1) PBS, 2) pac-Scr-PS (0.1 μmol/kg), 3) free AXL-PS (0.1 μmol/kg), and 4) pac-AXL-PS (0.1 μmol/kg). Samples were injected once every 3 days until day 39. The volume of tumors and weight of mice were recorded before every treatment and 3 days after the last treatment. Antitumor activity was evaluated in terms of tumor size by measuring two orthogonal diameters at various time points (V = 0.5 × LW^2^; L: length, W, width).

### Associated Content

Materials, experimental methods, synthesis and characterization of poly(serinol phosphodiester) bottlebrush polymer, additional experimental data including sequences, RP-HPLC chromatograms, MST binding affinity analysis, cell proliferation rate, western blot imaging, cell migration ratio, and live mice imaging.

## Author Contributions

K.Z. and T.S. devised the experiments and wrote the manuscript. T.S. conducted the synthesis of materials, purification, and material/biological characterization. All other authors contributed to material synthesis, purification, and/or discussion of the results. All authors edited the manuscript.

## Notes

K.Z. holds financial interest in pacDNA Inc., a company commercializing the pacDNA technology.

## Acknowledgments

The authors thank Dr. Guoxin Rong from the Institute for Chemical Imaging of Living Systems at Northeastern University for assistance with flow cytomery, Dr. Ambika Bajpayee for microscale thermophoresis analysis and Dr. Eno E. Ebong for western blot imaging analysis.

