## Supporting info for "Making Aptamers More Antibody-like: Targeting AXL *in Vivo* Using a Bottlebrush Polymer-Conjugated Aptamer"

**Material**

Phosphoramidites and supplies for DNA synthesis were purchased from Glen Research Co. (Sterling, VA, USA). Methoxy polyethylene glycol (PEG) glutaramide succinimidyl ester (M_n_=10 kDa) was purchased from Creative PEGWorks (Chapel hill, NC, USA). Human NCI-H1299 lung cancer cell line, human A549 lung cancer cell line, human LN229 glioblastoma cancer cell line and human SKOV3 ovarian cancer cell line were purchased from American Type Culture Collection (Rockville, MD, USA). All other materials were purchased from Fisher Scientific INC. (USA), Sigma-Aldrich Co. (USA), or VWR International LLC. (USA) and used as received unless otherwise indicated.

**Oligonucleotides and Quasar backbone synthesis^1^**

All oligonucleotides with or without poly(serinol phosphodiester) backbones were synthesized via solid-phase synthesis method on a Model 391 DNA synthesizer (Applied Biosystems, Inc., Foster City, CA). All natural/modified DNA phosphoramidites and serinol phosphoramidite were purchased from Glen Research Co. (Sterling, VA, USA). All oligonucleotide strands were cleaved from the CPG column and deprotected in aqueous ammonium hydroxide solution (28-30% NH_3_) at room temperature for 24 hours. The oligonucleotide with poly (serinol phosphodiester) backbones containing Fmoc group were deprotected on-column in DMF with 20% piperidine 3 times and washed with DMF 2 times. The CPG was dried in vacuo and cleaved via the same method as normal oligonucleotide strands. All Quasar backbones and oligonucleotide strands were purified by RP-HPLC, followed by the removal of dimethoxy-trityl (DMT) groups using 20% acetic acid and extracted 3 times with ethyl acetate.

**Transmission electron microscopy (TEM)**

Samples (10 μM) were placed on parafilm as a droplet, onto which a copper-coated TEM grid was gently placed. The grids were then moved, dried, and stained using 2% uranyl acetate for 10 min. TEM images were collected on a JEOL JEM 1010 electron microscope with an accelerating voltage of 80 kV.

**Dynamic light scattering (DLS)**

The samples were prepared by dissolving in Nanopure™ water at a concentration of 1 μM and filtering through a 0.2 μm PTFE filter prior to measurement. DLS measurements were conducted using a Malvern Zetasizer Nano-ZSP (Malvern, UK).

**N,N-dimethylformamide gel permeation chromatography (DMF-GPC)**

HPLC-grade DMF containing 0.05 M lithium bromide was utilized as the mobile phase, with sample analysis conducted at a flow rate of 0.4 mL/min. Calibration for DMF-GPC was performed using a ReadyCal kit of polyethylene glycol standards (PSS-Polymer Standard Service-USA Inc., MA, USA), covering an Mn range from 232 Da to 1015 kDa. Sample analysis was carried out on a Tosoh EcoSEC HLC-8320 GPC system (Tokyo, Japan) equipped with a TSKGel α-M 7.8×300 mm, 13 μm column and RI/UV-Vis detectors.

**Scheme S1. Synthesis of serinol-phosphoramidite (3)**

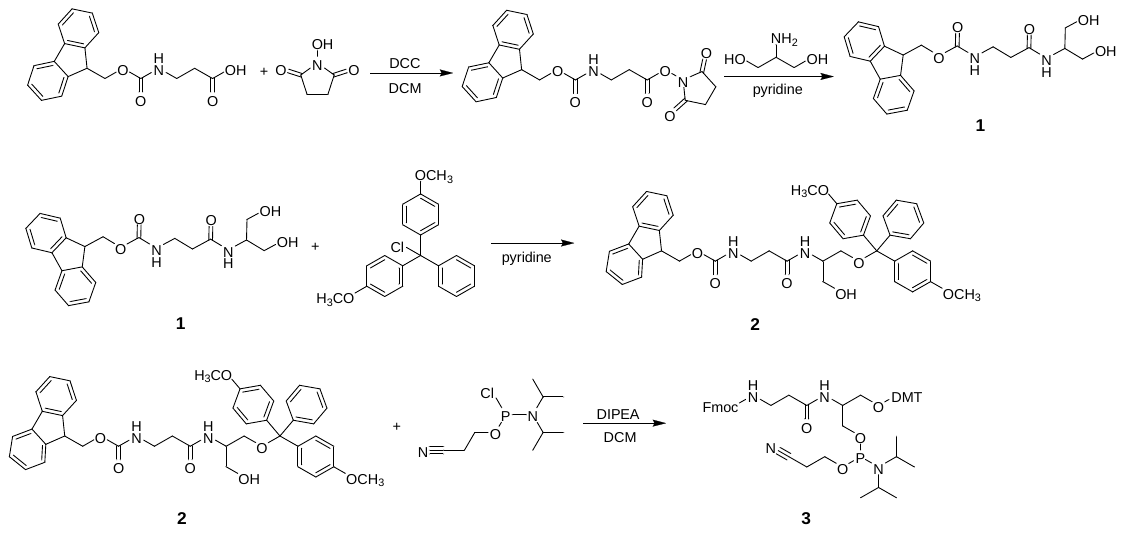

**Synthesis of Fmoc-β-Ala-serinol (1, Scheme S1)**

Fmoc-β-alanine (9.33 g, 30 mmol) and *N*-hydroxysuccinimide (3.45 g, 30 mmol) were dissolved in a mixed solvent of dichloromethane (DCM, 120 mL) and *N,N*-dimethylformamide (DMF, 6 mL). *N,N’*-dicyclohexylcarbodiimide (DCC, 6.18 g, 30 mmol) was dissolved in DCM (12 mL) and added to the mixture. The mixture was stirred at room temperature for 1.5 h, and a white precipitate of *N,N’*-dicyclohexylurea (DCU) was observed. The reaction mixture was filtered, and the liquid was transferred to a mixture containing serinol (2.73 g, 30 mmol) and pyridine (42 mL). The reaction mixture was allowed to stir for 16 h at room temperature, before the solvent was removed by evaporation to yield a semi-solid residue (containing residue pyridine). Toluene (50 mL) was added to the residue and were co-evaporated under reduced pressure 3× to remove pyridine. The resulting white solid was refluxed in DCM (200 mL) for 2 h, before being chilled to -20 °C. The product was then collected by filtration and washed with DCM (100 mL) 2× and diethyl ether (100 mL) 1×.^1^ The final product was dried under high vacuum and stored at -20 °C.

**Synthesis of Fmoc-β-ala-serinol-DMT (2)**

Fmoc-β-Ala-serinol (**1**, 5.88 g, 15.3 mmol) was placed in a flask filled with N_2_ and was dissolved in pyridine (16 mL). The flask was chilled in an ice bath. 4,4’-Dimethoxytrityl chloride (5.37 g, 15.9 mmol) was dissolved in pyridine (32 mL) and added dropwise to the mixture containing **1**. The mixture was allowed to stir under N_2_ for 1 h in ice bath and then overnight at room temperature. Methanol (1 mL) was added to the mixture and stirred for 15 min to quench the reaction. Removal of solvent under reduced pressure yields an oily residue, which was then co-evaporated with toluene (40 mL) 3×. The mixture was dissolved in DCM (50 mL) and washed with 5% sodium bicarbonate solution (50 mL) and then with brine (50 mL), before being dried over anhydrous sodium sulfate. The crude product was concentrated and purified by silica gel column purification (ethyl acetate/methanol/triethylamine 95:5:1 v:v:v).

**Synthesis of serinol-phosphoramidite (3)**

Fmoc-β-Ala-serinol-DMT (**2**, 2.76 g, 4 mmol) was placed in a flask charged with N_2_ and dissolved in dry DCM (12 mL) containing *N,N*-diisopropylethylamine (DIPEA, 3.5 mL). The flask was chilled in an ice bath. 2-Cyanothyl-*N,N*-diisopropylchlorophosphoramidite (1.9 g, 8 mmol) was dissolved in DCM (4 mL) and added dropwise to the mixture containing **2**. The reaction mixture was allowed to stir vigorously for 20 min before being warmed to room temperature and stirred for another 40 min. An excess of ethyl acetate was added to the reaction mixture. The mixture was washed with saturated sodium bicarbonate solution (10 mL). After drying over anhydrous sodium sulfate, the mixture was filtered, and the filtrate was concentrated for silica gel column purification (hexane/ethyl acetate/triethylamine 67:33:1 v:v:v).

**Synthesis of Quasar**

Quasar backbone (100 nmol) and NHS-terminated 10 kDa mPEG (1:1 amine:NHS ester) were dissolved in 1 mL of phosphate-buffered saline (PBS, pH 7.4). The mixture was shaken at 4 °C overnight and lyophilized to give a white powder, which was then redissolved in 1 mL anhydrous DMF containing 42 μL triethylamine. To this mixture, 1 equiv of NHS-terminated PEG (dissolved in 1 mL DMF) was added in 5 aliquots (12 h between each aliquot), and the mixture was gently shaken at room temperature. The mixture was dried *in vacuo* and purified by aqueous GPC.

**Table S1. Sequences used in this study.^2^**

| **Sample** | **Sequence** |
| --- | --- |
| PO | 5’-ATGACAATCGCCTCAATTCGACAGAGGCTCAC-3’ |
| PS | 5’-aUgaCAAUCGCCUCAaUUCGACAgGAGGCUCaC-3’ |
| SCR | 5’-GtGcTAATCCtACAATTcACAGCACaTGCGCgag-3’ |
| pac-AXL-PO | 5’-ATGACAATCGCCTCAATTCGACAGAGGCTCAC-(Ser)_30_- ATGACAATCGCCTCAATTCGACAGAGGCTCAC-3’ |
| pac-AXL-PS | 5’-aUgaCAAUCGCCUCAaUUCGACAgGAGGCUCaC-(Ser)_30_- aUgaCAAUCGCCUCAaUUCGACAgGAGGCUCaC-3’ |
| pac-Scr-PS | 5’-GtGcTAATCCtACAATTcACAGCACaTGCGCgag-(Ser)_30_-GtGcTAATCCtACAATTcACAGCACaTGCGCgag-3’ |
| Serinol Brush | 5’-(Ser)_30_-3’ |

Note. C,U = 2’-F-dN; a, g, c, t = 3’-monothio-dN; C,U = 2’-F-3’-monothio-dN;

A,C,G,T = naturel nucleotides; Ser = amine-serinol phosphoramidite

**Table S2. Plasma pharmacokinetics calculated parameters of 5’-Cy5-labeled aptamers, PSP pacDNAs and bottlebrush polymer in C57BL/6 mice.**

| **Sample ID** | **t_1/2α_ (h)** | **t_1/2β_ (h)** | **AUC_∞_ (nmol/mL·h)** |
| --- | --- | --- | --- |
| AXL-PO | 0.14 | 3.88 | 5.91 |
| pac-AXL-PO | 2.53 | 25.59 | 84.71 |
| AXL-PS | 0.48 | 3.62 | 9.361 |
| pac-AXL-PS | 2.37 | 29.64 | 66.12 |
| Brush | 1.50 | 30.99 | 271.9 |

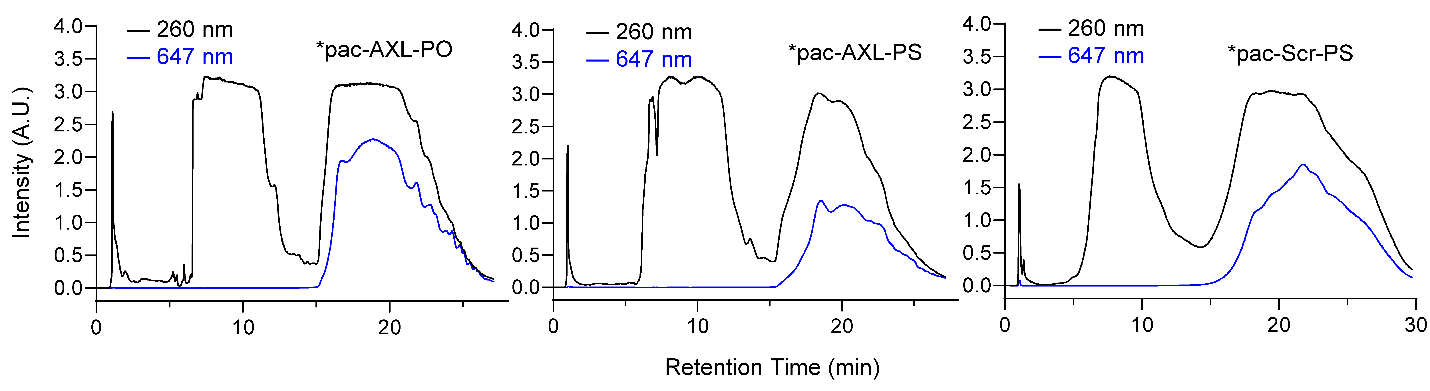

**Figure S1.** RP-HPLC chromatograms of PSP pacDNA backbones. The asterisk-labeled peaks were collected for further reaction.

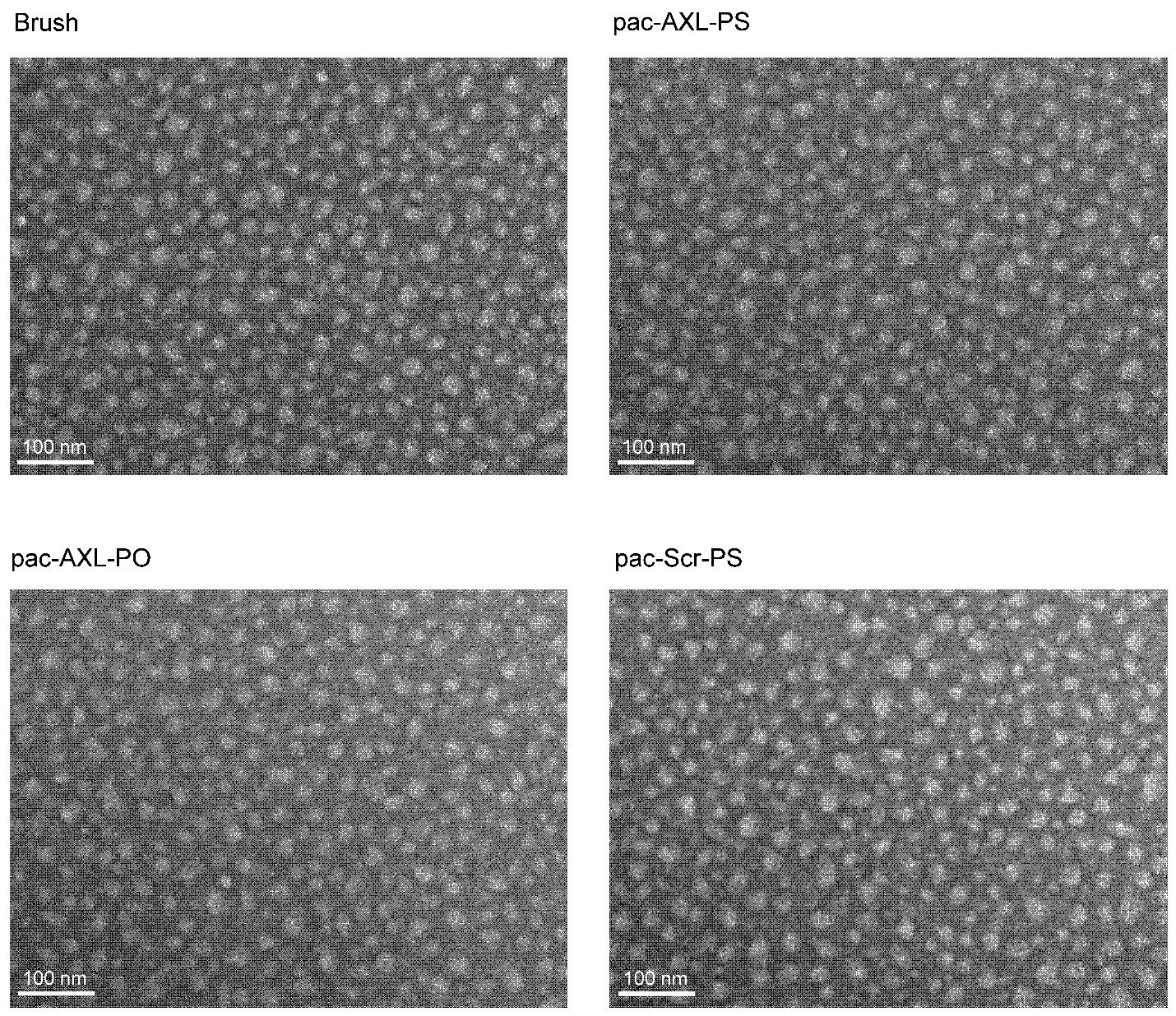

**Figure S2.** TEM images (negative staining by 2% uranyl acetate) of bottlebrush polymer, pac-AXL-PS, pac-AXL-PO, and pac-Scr-PS.

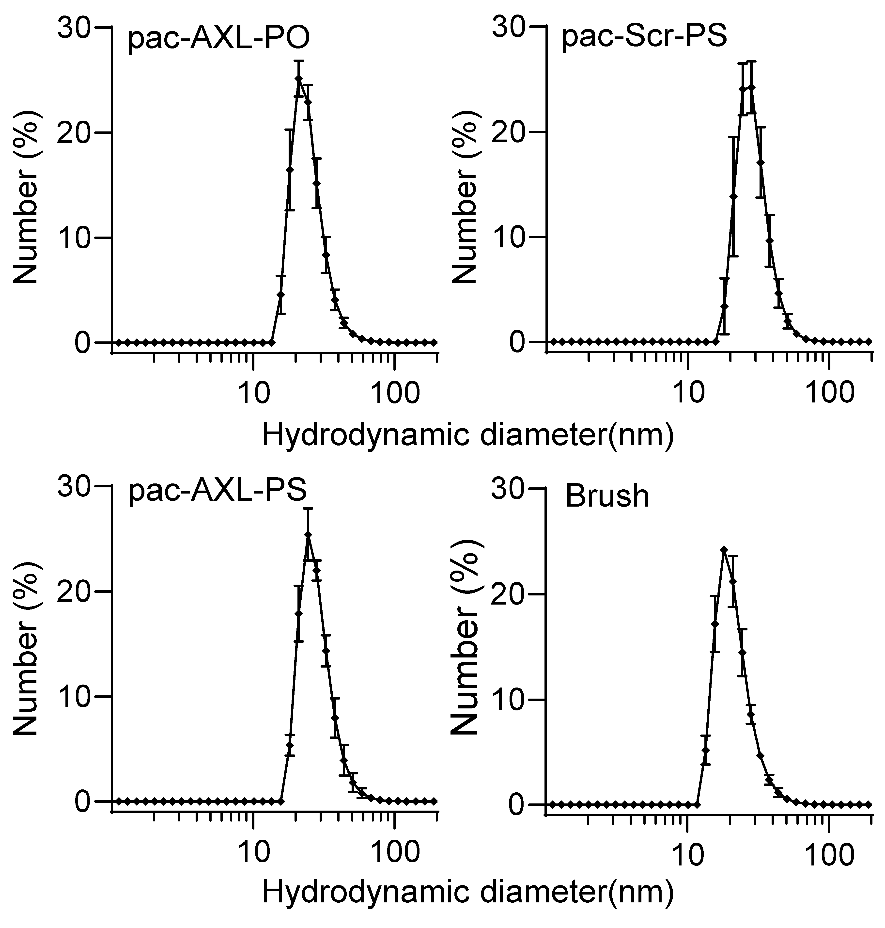

| **Sample ID** | Pac-AXL-PO | Pac-AXL-PS | Pac-Scr-PS | Brush |
| --- | --- | --- | --- | --- |
| **Hydrodynamic diameter(nm)** | 24.3±7.3 | 27.7±8.6 | 31.0±9.1 | 21.6±6.3 |

**Figure S3.** DLS measurements of PSP bottlebrushes, pac-AXL-PO, pac-Scr-PS, pac-AXL-PS in Nanopure™ water.

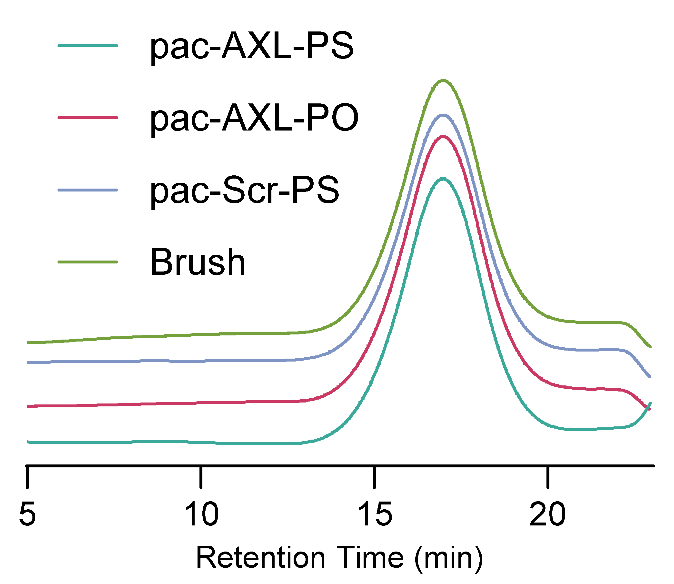

| **Sample ID** | **M_n_(kDa)** | **M_w_(kDa)** | **PDI** |
| --- | --- | --- | --- |
| pac-AXL-PO | 317.7 | 467.1 | 1.47 |
| pac-AXL-PS | 301.8 | 362.6 | 1.20 |
| pac-Scr-PS | 302.2 | 352.3 | 1.16 |
| Brush | 298.4 | 389.2 | 1.34 |

**Figure S4.** DMF-GPC chromatogram and average molecular weight of pacDNAs and bottlebrush polymer

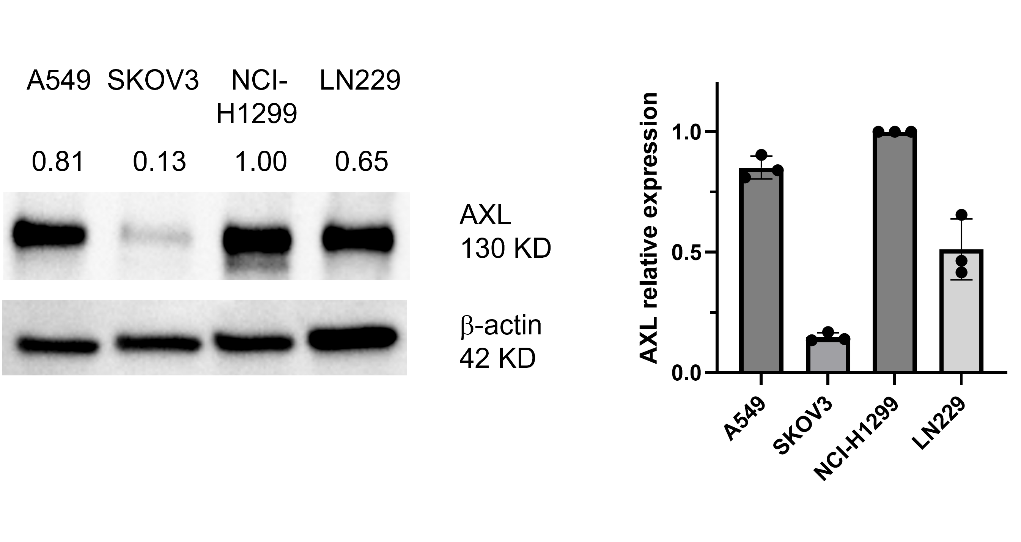

**Figure S5.** AXL expression level of LN229, SKOV3, NCI-H1299 and A549 cell lines determined by immunoblotting.

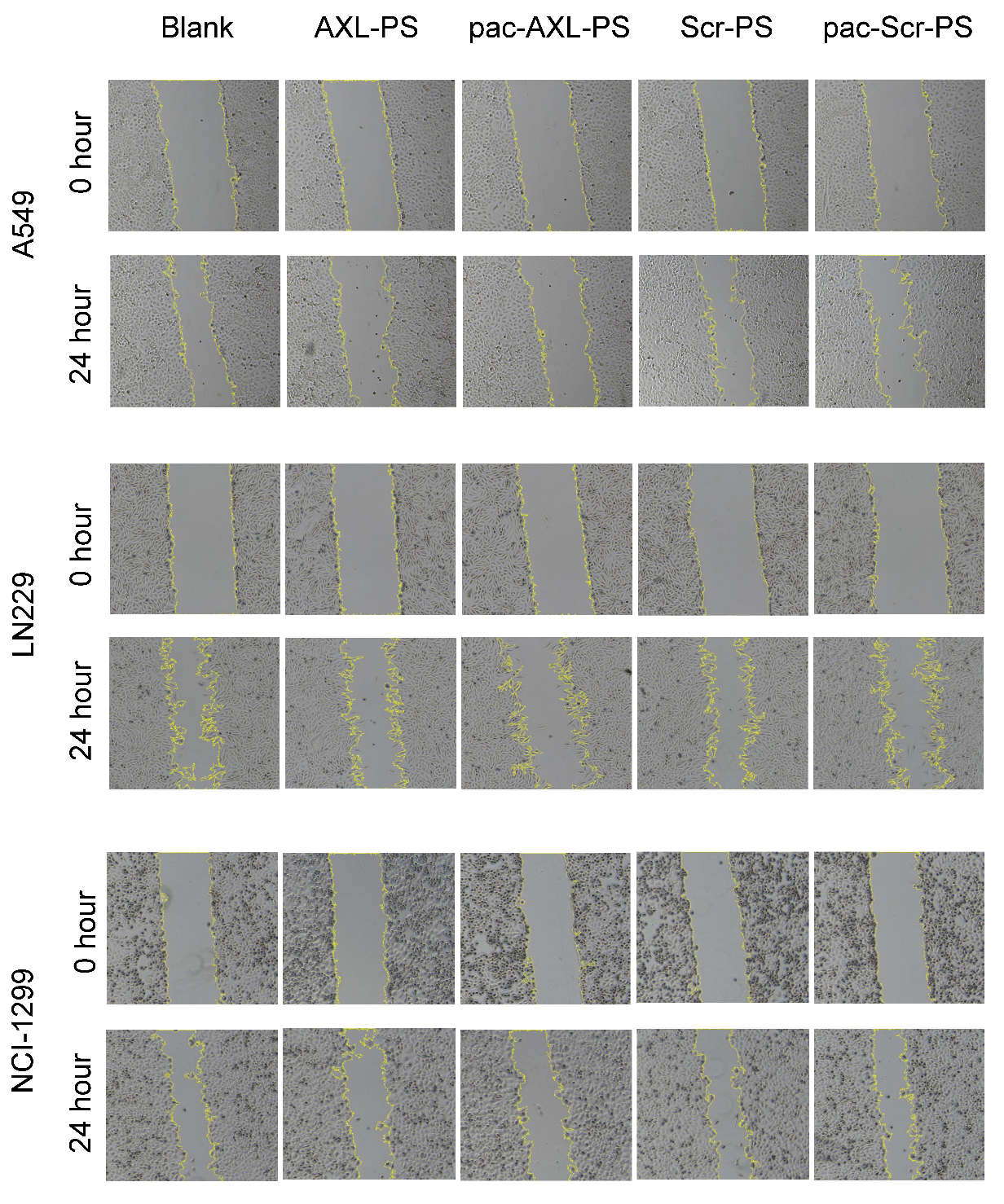

**Figure S6.** Wound healing assays were performed to assess A549, LN229 and NCI-H1299 cell migration. The healing area was determined 24 h after the scratch.

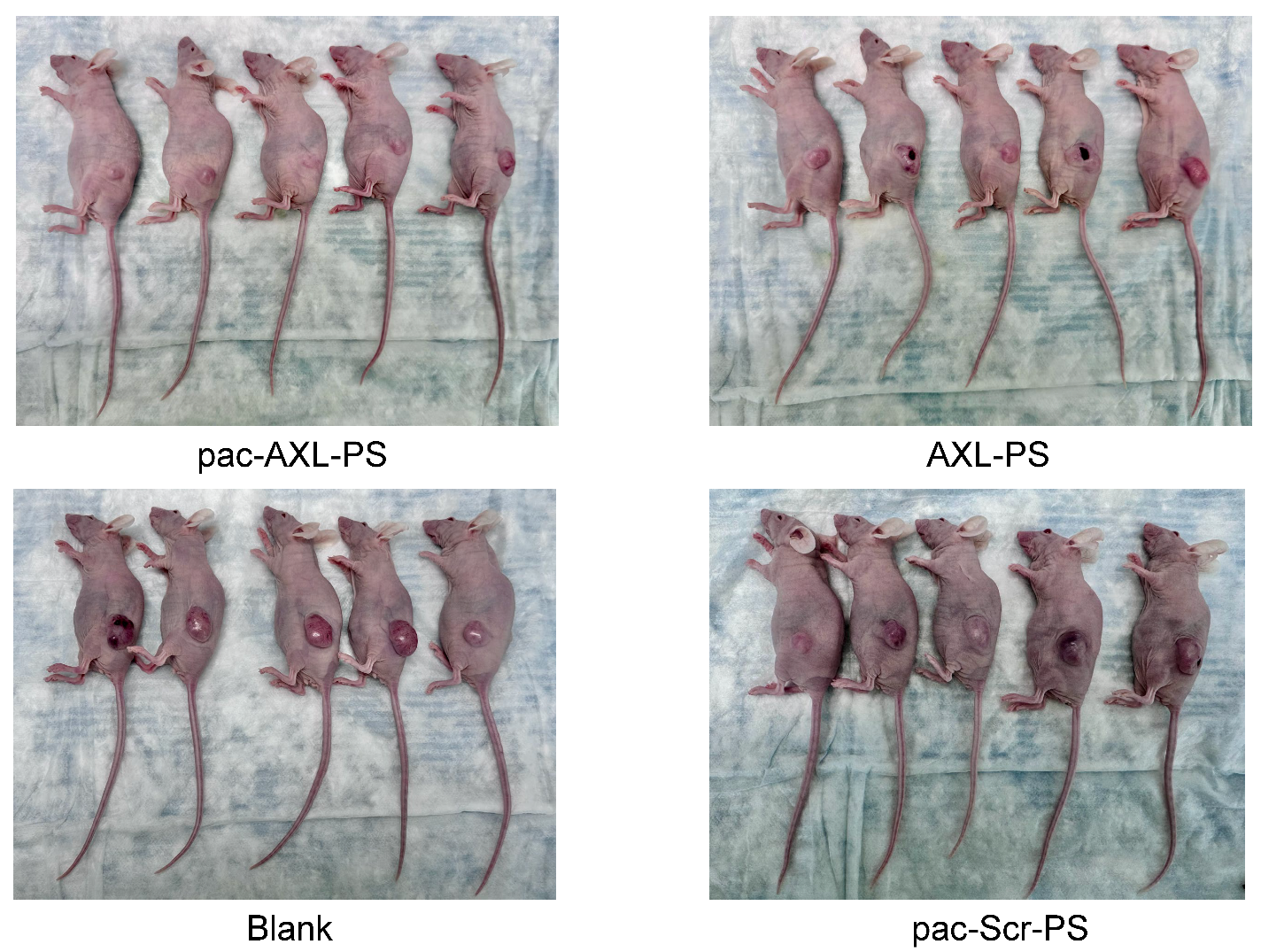

**Figure S7.** The tumor growth of each mouse was monitored and representative tumors on control and treated mice from four groups after 31 days of treatment are presented.
